# Short-term monocular deprivation boosts neural responsiveness to audio-visual events for the undeprived eye

**DOI:** 10.1101/2022.05.17.492319

**Authors:** A. Federici, G. Bernardi, I. Senna, M. Fantoni, M.O. Ernst, E. Ricciardi, D. Bottari

## Abstract

A brief period of monocular deprivation (MD) induces short-term plasticity of the adult visual system. Whether MD elicits changes beyond visual processing is yet unclear. Here, we assessed the specific impact of MD on multisensory processes. Neural oscillations associated with visual and audio-visual processing were measured for both the deprived and the undeprived eye. Results revealed that MD changed neural activities associated with unimodal and multisensory processes in an eye-specific manner. Selectively for the deprived eye, alpha activity was reduced within the first 150 ms of visual processing. Conversely, gamma activity was enhanced in response to audio-visual events only for the undeprived eye within 100-300 ms after stimulus onset. The analysis of gamma responses to unimodal auditory events revealed that MD elicited a crossmodal upweighting for the undeprived eye. Distributed source modeling suggested that the right parietal cortex played a major role in all neural effects induced by MD. Finally, visual and audio-visual processing alterations emerged selectively for the induced (but not the evoked) component of the neural oscillations, indicating a major role of feedback connectivity. These findings support a model in which MD increases excitability to visual events for the deprived eye and to audio-visual and auditory events for the undeprived eye. On the one hand, these results reveal the causal impact of MD on both unisensory and multisensory processes but with distinct frequency-specific profiles. On the other hand, they highlight the feedback nature of short-term neural plasticity. Overall this study shed light on the high flexibility and interdependence of unimodal and multisensory functions.

**Highlights:**

- We unveiled the impact of temporary MD on visual and audio-visual processing
- MD enhanced visual excitability for the deprived eye
- MD boosted neural responses to audio-visual events for the undeprived eye
- Analyses of auditory processing revealed crossmodal effects following MD
- Short-term MD primarily affects induced, non-phase-locked, oscillatory activity

## Introduction

While neuroplasticity of the visual system is maximal during development (Hubel and Wiesel, 1970; Hensch, 2005; Espinosa and Stryker, 2012; Levelt and □ubener, 2012; Reh et al., 2020), evidence that a certain degree of plasticity is retained in the adult visual cortex has accumulated (Karmarkar and Dan, 2006; Spolidoro et al., 2009; Espinosa and Stryker, 2012; Hensch and Quinlan, 2018). Studies in adults employing psychophysics and neuroimaging methods showed that a brief period of MD, a pivotal model for testing the plasticity of the visual system, alters the ocular balance by strengthening visual processing of the deprived eye and weakening the undeprived eye (Castaldi et al., 2020). These functional changes reflect ocular dominance shifts in V1 in favor of the deprived eye (Lunghi et al., 2015a; Binda et al., 2018) and are supposedly driven by homeostatic plasticity (Lunghi et al., 2015b), the mechanism involved in preserving the cortical excitatory-inhibitory balance (Turrigiano, 2012). Besides this well-documented MD effect on visual processing, little is known about how MD affects multimodal processing.

Alterations induced by MD seem to impact beyond visual processing. At the behavioral level, short-term MD was found to affect the interplay between sensory modalities (Lo Verde et al., 2017; Opoku-Baah and Wallace, 2020). Overall, results were consistent with a decreased multisensory interplay for the deprived eye, in which the visual processing is generally strengthened following MD, and vice versa, an increased multisensory interplay for the undeprived eye, in which visual processing is typically weakened following MD. However, the neural correlates of MD effects on multisensory processes are still unknown. To fill this gap, we exploited the model of MD to elicit short-term plasticity and investigated specific changes in visual and audio-visual processes for the deprived and undeprived eye. Overall, this study aimed to address whether a brief period of monocular visual experience in adulthood can affect the neural processing associated with multisensory audio-visual events, and which are the underpinning neural mechanisms.

Here, we combined psychophysical and electrophysiological approaches. We employed the *sound-induced flash illusion* (Shams et al., 2000, Hirst et al., 2020), in which the number of perceived flashes can be biased by the number of concurring beeps. To investigate the impact of MD on visual processing, we measured neural oscillations in response to a single flash before and after a deprivation phase. To determine the existence of an audio-visual MD effect, the analysis focused on neural oscillations elicited by the *fission* illusion trials in which a single flash is coupled with two beeps, and two flashes can be perceived (Shams et al., 2000). This illusion reveals the impact of auditory input on visual processing in case of conflictual audio-visual information.

Neural correlates associated with visual and audio-visual processing are characterized by their spectral, temporal, and spatial properties. Reflecting high and low neuronal excitability cycles, neural oscillations reveal perceptual organization in both visual (Jensen et al., 2014; VanRullen, 2016) and multisensory analyses (Cooke et al., 2019; Lennert et al., 2021). We expected MD to induce changes in alpha activity, which is known to reflect levels of inhibition (Klimesch et al., 2007; Klimesch, 2012), and, thus, increase the visual system excitability for the deprived eye (Lunghi et al., 2015a). Moreover, we hypothesized MD to alter responses in the gamma range. Gamma activity has been as well associated with excitatory-inhibitory balance (Jensen et al., 2010; 2012), and its modulation was found to be involved in eliciting a perceptual fission illusion (Bhattacharya et al., 2002; Mishra et al., 2007; Lange et al., 2011, 2013; Balz et al., 2016).

Importantly, distinct components of neural oscillations characterize different types of processing according to the direction of information flow: while the evoked activity, which is phase-locked to the stimulation, has mainly been associated with feedforward processing (thalamo-cortical), the induced oscillatory activity, which is not phase-locked to the onset of the stimulus, has mainly been associated with feedback processing (cortico-cortical connectivity, Klimesch et al., 1998; Tallon-Baudry and Bertrand, 1999; Chen et al., 2012). Studies in humans and non-human animal models demonstrated that permanent sensory deprivation primarily affects induced oscillatory activity (Bottari et al., 2016; Yusuf et al., 2017; Bednaya et al., 2021). As homeostatic plasticity is an intrinsic feedback mechanism (Turrigiano and Nelson, 2004), we predicted to primarily observe changes in induced oscillatory activity following temporary MD for both visual and audio-visual processing. Finally, source modeling was employed to investigate the neural sources of short-term plasticity effects.

## Materials and Methods

### Participants

Since the effect of MD on multisensory processing was unknown, we estimated the minimum sample size needed to reach the expected effect of MD on visual processing as previously reported in the literature. We expected the MD effect on visual processing to be at occipito-parietal electrodes, in the alpha range [8-14 Hz] (Lunghi et al., 2015a), and within the first stages of visual processing [0-120 ms] (comprising the earliest visual evoked potential, C1 wave, known to be modulated by MD; Lunghi et al., 2015a). The power analysis was performed by simulating our planned analysis (t1-t0 Deprived eye vs. t1-t0 Undeprived eye) on the alpha frequency power using a cluster-based permutation test (Wang and Zhang, 2021). The analysis revealed an estimated minimum sample size of 17 participants (for further details on sample size estimation see Supplementary Materials and Figure S1). Note that previous studies investigating the effect of MD using EEG analyzed up to 16 participants (Lunghi et al., 2015a; Zhou et al., 2015; Schwenk et al., 2020).

To determine individual suitability for the *main experiment* (EEG experiment), twenty-seven potential participants completed a *preliminary behavioral assessment* (see the section below) in order to assess whether they met the following inclusion criteria: (i) to perceive the *fission illusion* with the dominant eye (i.e., >20% illusory rate), and (ii) to not be completely biased by the sound in the illusory conditions (i.e., >95% illusory rate). Out of twenty-seven young adults tested (mean age 28.22 ± 2.41 SD, twelve males and fifteen females), six participants were excluded as they did not meet these inclusion criteria or could not comply with the experimental instructions (see Supplementary Materials). Out of the 21 participants who performed the *main experiment*, one further participant was excluded due to his poor behavioral performance (the number of errors was 3 SD above the group mean in the conditions in which only auditory stimuli were presented, i.e., the control conditions). The final sample included 20 young-adult participants (mean age 28.45 ± 2.67 SD, eight males and twelve females). They all had normal or corrected-to-normal vision (visual acuity ≥ 8/10) and did not report hearing deficits or a history of neurological conditions. Since one EEG and one behavioral dataset, from different participants, went lost due to technical issues during acquisitions, the analyzed data sample included 19 behavioral and 19 EEG datasets.

The study was approved by the local Ethical Committee (Comitato Etico di Area Vasta Nord Ovest Regione Toscana protocol n. 24579). Each participant signed a written informed consent before taking part in the experiment. The experimental protocol adhered to the principles of the Declaration of Helsinki (2013).

### Stimuli and apparatus

The experiment was performed in a dimly lit and sound-attenuated chamber (BOXY, B-Beng s.r.l., Italy). Participants were comfortably sitting in front of the apparatus, with their eyes at a distance of 60 cm from the monitor. Visual stimuli were presented on an LCD monitor (60 Hz refresh rate; 24.5 inches; 1920 × 1080 screen resolution), and audio stimuli were delivered via a single speaker (Bose® Companion 2, series III multimedia) located below the screen and aligned with its center. Stimuli were flashes and beeps. Both visual and audio stimuli were created using Matlab (The Mathworks, Inc. - version 2017b). The audio stimulus was a 7 ms quadratic beep with a 3.5 kHz frequency and a sampling rate of 44.1 kHz, which was presented at about 75 dB. The visual stimulus was a 2° diameter grey dot displayed 5° below the center of the screen for 17 ms (corresponding to 1 frame) on a black background. The contrast level of the grey dot was selected individually via a staircase procedure to elicit the fission illusion in about 50% of trials (Pérez-Bellido et al., 2015, see below *Preliminary behavioral assessment*). Stimuli were delivered using E-Prime® software (version 2, Psychology Software Tools, Inc. www.pstnet.com). The Audio/Visual (AV) Device (Electrical Geodesics, Inc.) was employed to ensure accurate optimal synchronization between the presented stimuli and the recorded EEG traces.

### Experimental Design

The whole procedure consisted of two main parts performed on two separate days: a *preliminary behavioral assessment* and the *main experiment*, in which the dominant eye was deprived of patterned visual input using a translucent eye patch. In both the *preliminary behavioral assessment* and the *main experiment*, participants performed a *monocular visual discrimination task*.

#### Monocular visual discrimination task

Participants were asked to report the number of perceived flashes (0, 1, or 2) while task-irrelevant beeps (0, 1, or 2) were presented. Responses were given by pressing one of three keypad buttons using the right-hand fingers. In each trial, audio (A) and visual (V) stimuli could be presented coupled or isolated constituting eight conditions: half the conditions were unisensory and comprised single or couples of visual or auditory events (visual: V and VV; auditory: A and AA) and the other half were multisensory (coherent audio-visual stimulation: AV and AVAV; illusory audio-visual stimulation: AVA and VAV, for a schematic summary of all conditions see the table reported in Figure 1a). Unisensory auditory trials (i.e., A and AA) represented control conditions. They were employed to ensure that participants correctly performed the task (participants with a number of errors 3 SD above the group mean were excluded from the analyses as described in the Participants section). The presentation order of the conditions was randomized.

**Figure 1.**
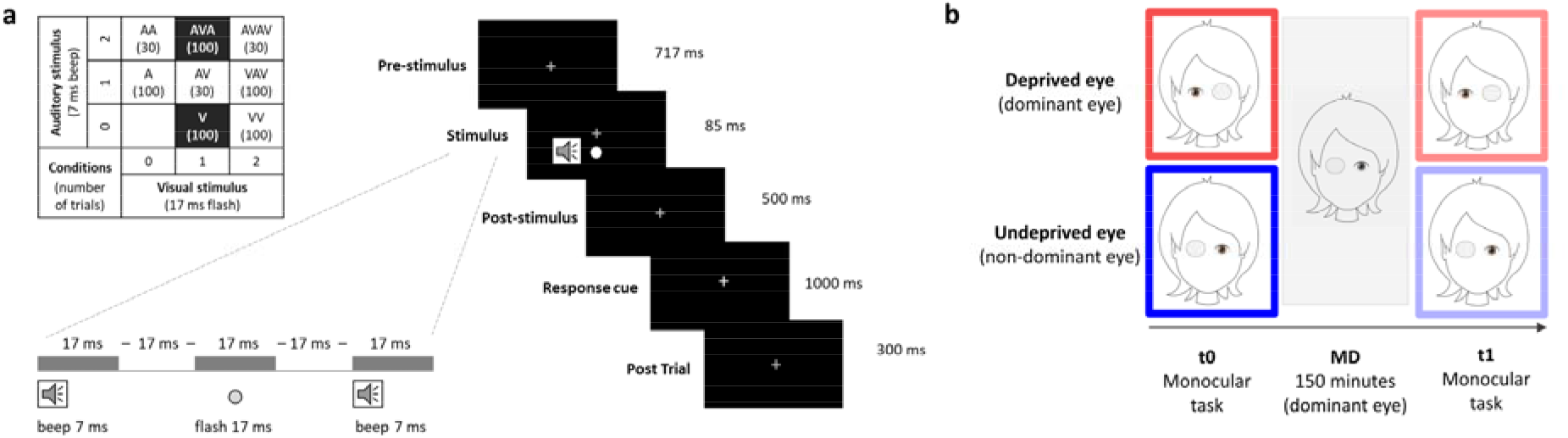
Schematic illustration of the experimental paradigm. (a) The top left table shows all the possible combinations of the stimuli (visual stimuli V and auditory stimuli A), representing all the eight possible conditions (conditions of interest are highlighted, and the number of trials in each condition is shown in brackets). On the right, an example of a single trial presentation in AVA condition is depicted. (b) Experimental procedure showing the four sessions performed with the Deprived eye (upper line, red contour) and the Undeprived eye (bottom line, blue contour). At t0 and t1, participants performed the visual monocular task with each eye (eye order was counterbalanced across participants). Between t0 and t1, participants wore a translucent eye patch on the dominant eye (Deprived eye) for 150 minutes (MD phase).

Since our main aim was to explore changes in the neural response to audio-visual events caused by a short-term MD, the main analysis focused on the audio-visual condition inducing fission illusion (AVA). Generating an unstable percept, this illusion allows investigating subtle changes in audio-visual processing. As a control, we investigated changes in the unisensory visual condition (V). Notably, the visual stimulus was the same single flash in AVA and V conditions.

All trials started with a grey fixation cross presented at the center of the screen on a black background. After 717 ms, the stimulation was delivered. In the case of the fission illusion (i.e., AVA), each visual and auditory stimuli were separated by one frame (i.e., 17 ms; see an example for AVA condition in Figure 1a and Supplementary Materials). Following the stimulation, a blank screen appeared for 500 ms (response-free time window), then the fixation cross became white, and participants were asked to respond within 1 second (except for the staircase procedure, in which participants had infinite time to respond; see *Preliminary behavioral assessment*). As soon as the response was given, a blank screen was presented for 300 ms before the beginning of the subsequent trial. Participants were asked to maintain their gaze at the fixation cross throughout the duration of the trial.

#### Preliminary behavioral assessment

The ocular dominance via the Porta Test (see Supplementary Materials), the visual acuity via the eye chart, and the rate of the illusory percept during the *monocular visual discrimination task* were measured to verify if the participant could take part in the *main experiment*.

Once ocular dominance and visual acuity were tested, participants performed two short versions of the *monocular visual discrimination task*. The first short version comprised a staircase procedure (see Supplementary Materials for details) that was used to identify, at the individual level, the luminance contrast between the grey dot and the black background needed to elicit the fission illusion in about 50% of trials (Pérez-Bellido et al., 2015). Participants performed this test monocularly, with the dominant and non-dominant eye (the order was randomized). The contrast level for the dominant and non-dominant eye did not differ within-participant (t(26)= −1.466, p=0.155). The second short version of the *monocular visual discrimination task* (30 trials for each of the following conditions V, VV, AVA, VAV, A, and 6 for AA, AV, AVAV) was then performed with the dominant eye (with the contrast identified by the staircase procedure) to evaluate whether the participant was fulfilling the inclusion criteria to take part in the *main experiment* (see Participants section).

Previous evidence revealed that individual sensory preference (*Audio* or *Visual*) impacts on multisensory processing (Giard and Perronet, 1999). To assess whether individual sensory predisposition might affect multisensory short-term plasticity, participants monocularly performed a speeded object recognition task based on auditory, visual, and audio-visual information (see Supplementary Materials for details). Each participant’s Sensory-Preference (*Audio* or *Visual*) was classified for each eye, and no significant difference was found between them (p>0.68). Sensory-Preference was an additional measure that we took into consideration: for this reason, the N within the *Audio* and *Visual* groups were not balanced.

#### Main experiment

Participants recruited in the *main experiment* came to the laboratory a second time on a different day. Since the *main experiment* lasted about five hours, the data were always acquired in the morning (approximately between 9 am and 2 pm) to avoid possible confounds associated with the circadian cycle or related to (visual) activities performed before the experiment. Each participant repeated the short version of the *monocular visual discrimination task* with the dominant eye comprising the staircase procedure to ensure the test-retest reliability of the selected visual stimulus contrast (no significant difference was found between the contrast levels measured in the two assessments; t(20)=0.637, p=0.532).

A brief practice of 16 trials was run before the *main experiment*. Participants performed the *monocular visual discrimination task*, with each eye, before (t0) and after (t1) a period of monocular deprivation (MD) (see Figure 1b), while their EEG signal was recorded. Thus, each participant performed a total of four sessions of the *monocular visual discrimination task* (i.e., at t0 and t1, both with the dominant and the non-dominant eye). Note that whether they started with the dominant or the non-dominant eye was counterbalanced across participants. MD consisted of 150 minutes in which the dominant eye was occluded by a translucent eye patch, following a validated procedure (Lunghi et al., 2011). From now on, we will refer to the dominant eye as the ‘Deprived eye’ and to the non-dominant eye as the ‘Undeprived eye’.

To prompt active and controlled multisensory interactions with the environment during the MD phase, participants were engaged in predefined activities. Since the physical activity was demonstrated to boost short-term homeostatic plasticity in the adult visual cortex (Lunghi and Sale, 2015), all participants were engaged in the following activities: table football, table hockey, ping pong, and billiards, each of them lasting 15 minutes. The EEG cap was always kept on the scalp for the entire experiment duration.

Each monocular session (i.e., t0 Deprived, t0 Undeprived, t1 Deprived, and t1 Undeprived) comprised 100 trials for the conditions V, VV, AVA, VAV, A, and only 30 trials for the conditions AA, AV, AVAV, and was divided into five blocks (118 trials each) lasting about 5 minutes each. The number of trials was chosen to keep the duration of the monocular session within the estimated length of the MD effect (which has been demonstrated to be present for up to 90 minutes, but substantially decrease after 15 minutes; see Lunghi et al., 2011).

### EEG recording and preprocessing

EEG data were collected continuously during the four monocular task sessions (i.e., t0 Deprived, t0 Undeprived, t1 Deprived, and t1 Undeprived), using Electrical Geodesics EEG system with 64-channels (EGI; 500 Hz sampling rate). Offline, the data of the four sessions were concatenated at the individual level to detect common stereotypical artifacts. Data were preprocessed by implementing a validated approach (Stropahl et al., 2018; Bottari et al., 2020). The continuous recordings were filtered (low-pass cut-off at 40 Hz, Hanning filter, order 500; high-pass cut-off at 1 Hz, Hanning filter, order 100) and downsampled to 250 Hz to reduce the computational time. The filtered and downsampled data were segmented into consecutive 1-second epochs and cleaned using joint probability criterion: segments displaying an activity with a joint probability across all channels larger than 3 SD were removed (pop_jointprob function of EEGLAB; Delorme et al., 2007). Independent Component Analysis (ICA) based on the extended Infomax (Bell and Sejnowski, 1995; Jung et al., 2000a, 2000b) was then performed. The resulting ICA weights were saved and applied to the raw continuous unfiltered data (Stropahl et al., 2018; Bottari et al., 2020). Components associated with stereotypical artifacts, such as eye blinks and eye movements, were identified and removed using a semiautomatic procedure (CORRMAP, Viola et al., 2009). The data were then low-pass and high-pass filtered (100 Hz, filter order 100; 0.1 Hz, filter order 500) with a Hanning filter. Noisy channels were identified based on visual inspection and then interpolated using spherical spline interpolation (mean interpolated electrodes per subject 2.32 ± 2.26 SD) and re-referenced to the average. Finally, the residual power line fluctuations at 50 Hz were removed using the CleanLine EEGLAB plugin (https://github.com/sccn/cleanline). The EEG data were then split again into the original four sessions. Each session was then segmented into epochs of 2.2 seconds, from −1 to 1.2 seconds with respect to the onset of the stimulation. Noisy epochs were then rejected based on the joint probability across channels (Delorme et al., 2007) with a threshold of 3 SD (mean epochs rejected per subject in each session for V condition: t0 Deprived 12% ± 4 SD, t0 Undeprived 12% ± 6 SD, t1 Deprived 14% ± 5 SD, and t1 Undeprived 12% ± 5 SD, and for AVA condition: t0 Deprived 12% ± 6 SD, t0 Undeprived 12% ± 5 SD, t1 Deprived 12% ± 5 SD, and t1 Undeprived 14% ± 6 SD). All these steps were performed with EEGLAB software (Delorme and Makeig, 2004). Data were then imported into Fieldtrip (Oostenveld et al., 2011) to perform time-frequency decomposition and statistical analyses.

### Time-frequency decomposition

Time-frequency decomposition of the EEG data was performed within each session and separately for the visual and audio-visual conditions. Within each condition and session, we first extracted the induced power at a single trial level after subtracting the evoked activity (that is, subtracting from each trial the ERP computed averaging across trials without low-pass filtering). Time-frequency decomposition of single-trials was computed at each channel, separately for low (2-30 Hz) and high (30-80 Hz) frequency ranges. The oscillations in low frequencies were estimated using a Hanning taper with a frequency-dependent window length (4 cycles per time window) in steps of 2 Hz. Oscillations with higher frequencies were estimated using a Multitapers method with Slepian sequence as tapers, in steps of 5 Hz with a fixed-length time window of 0.2 seconds and fixed spectral smoothing of ± 10 Hz. For both frequency ranges, the power was extracted over the entire epoch (from −1 to 1.2 seconds) in steps of 0.02 seconds. Then, the average across trials was computed at the individual level within each session (t0 Deprived, t0 Undeprived, t1 Deprived, t1 Undeprived), conditions (V and AVA), and frequency range (low and high). The resulting oscillatory activity was baseline-corrected to obtain the relative signal change with respect to the baseline interval. For the low-frequency range, the baseline was set between −0.7 and −0.3 seconds, while for the high-frequency range, it was between −0.2 and −0.1 seconds. The low-frequency range, having longer cycles, required a wide baseline for the appropriate estimation of slow oscillations and was kept temporally distant from the stimulus onset to avoid temporal leakage of post-stimulus activity into the baseline period. The same procedure, without ERP subtraction from single trials, was implemented to estimate the total power. The baseline-corrected evoked power was computed by subtracting the baseline-corrected induced power from the baseline-corrected total power.

### Source reconstruction

To better characterize the neural alterations induced by MD, source estimation of the neural effects was performed using Brainstorm software (Tadel et al., 2011) on preprocessed EEG data. Sources were extracted by applying a dynamic statistical parametric mapping (dSPM; Dale et al., 2000), adopting minimum-norm inverse maps to estimate the locations of scalp electrical activities. For each dataset, we used single trials pre-stimulus baseline [−0.1 to 0.002 s] to calculate single subject noise covariance matrices and to estimate individual noise standard deviations at each location (Hansen et al., 2010). The boundary element method (BEM) provided in OpenMEEG was adopted as a head model; the model was computed on the first dataset and then applied to all the others (default parameters in Brainstorm were selected). Source estimation was performed by selecting the option of constrained dipole orientations (Tadel et al., 2011). Time-frequency decomposition was computed for each participant on the estimated sources at the single-trial level using the same approach described for the time-frequency decomposition performed at the sensor level.

### Statistical analysis

#### Behavioral data

For each participant, we computed the d-prime (d’) as visual and audio-visual sensitivity indices: d’ = *z*(*p* hits) - *z*(*p* false alarms), where *z* is the inverse cumulative normal function, and *p* is the proportion of hits and of false alarms out of signal and noise, respectively. Values equal to 0 or 1 were corrected as 1/n and (n-1)/n, respectively, with n being the number of signal or noise trials. To compute the visual d’, we defined as hits, trials in which participants perceived one flash (V condition) and correctly responded ‘one’. Consequently, false alarms were trials in which two flashes were presented (VV), and participants responded to having seen one flash. To compute the audio-visual d’, we defined, coherently with the fission illusion literature, false alarms AVA trials in which participants reported two flashes (Watkins et al., 2006; Whittingham et al., 2014; Pérez-Bellido et al., 2015; Vanes et al., 2016; Keil, 2020). Thus, AVAV trials in which participants correctly responded ‘two flashes’ were considered hits. Note that the audio-visual sensitivity (d’) is inversely related to the amount of fission illusion: smaller d’ indicated a greater amount of illusory percepts and vice-versa (note that Response means of each condition are reported in Supplementary Materials Figure S2).

Two mixed-design ANOVA, with Eye (Deprived and Undeprived) and Time (t1 and t0) as within-subjects factors, and Sensory-Preference (*Audio* and *Visual*) in the Deprived eye and Sensory-Preference (*Audio* and *Visual*) in the Undeprived eye as between-subjects factors, were performed separately on visual and audio-visual d’ values. The Sensory-preference between-subjects factors were inserted in order to control their impact on the visual and audio-visual perception.

#### Neural oscillations

Oscillatory activity occurring after stimulus onset was analyzed separately for visual (V) and audio-visual (AVA) trials to assess MD impact on visual and audio-visual processing. Time-frequency analyses were separately performed for induced and evoked oscillatory activity for both low [4-30 Hz] and high [30-80 Hz] frequency ranges.

To assess the impact of MD, we subtracted the oscillatory activity recorded before MD from the oscillatory activity recorded after MD (i.e., t1 minus t0). This difference was computed separately for the Deprived and Undeprived eye. From now on, *PowChangeDeprived* represents relative changes in power due to MD for the Deprived eye, and *PowChangeUndeprived* the relative changes in power due to MD for the Undeprived eye.

To compare the impact of MD on Deprived and Undeprived eye, a series of non-parametric cluster-based permutation tests were performed via a paired-sample t-test without a priori assumptions (i.e., across all electrodes, time-points, and frequencies) between *PowChangeDeprived* and *PowChangeUndeprived*. We used the Monte Carlo method with 1000 random permutations; cluster-level statistics were calculated taking the sum of the t-values within every cluster, with an alpha level of 0.05 (two-tailed) and a minimum neighbor channel = 1. Identified clusters were considered significant at p<0.025 (corresponding to a critical alpha level of 0.05 in a two-tailed test). We focused on the post-stimulus activity, and thus, statistical tests were performed for the entire response-free time window, that is, from 0 to 0.5 seconds. The time period after 0.5 seconds from the stimulation onset was discarded to prevent including motor artifacts. If a significant difference due to MD emerged in this test (*PowChangeDeprived* vs. *PowChangeUndeprived*), we assessed whether differences between the two eyes emerged at t0 or t1. To this end, two planned comparisons (i.e., t0 Deprived vs. t0 Undeprived and t1 Deprived vs. t1 Undeprived) were performed using the same cluster-based permutation analysis approach and the same parameters reported above. In case a significant difference between Deprived and Undeprived eye would emerge only at t1 and not at t0, it would be indicative of a specific effect of the MD manipulation and rule out possible differences between the two eyes at baseline.

#### Correlations between neural and behavioral changes

After assessing the normality of the data with Shapiro-Wilk tests, Pearson correlations were employed to assess whether the neural changes related to short-term MD were associated with behavioral changes. When multiple correlations were performed to the same dataset, their results were compared employing a bootstrap method adapted for independent samples (see https://github.com/GRousselet/blog/tree/master/comp2dcorr).

The datasets and code used in the present study are available from the corresponding author on reasonable request.

## Results

### Behavioral data

#### Unisensory visual

The mixed-design ANOVA on visual d’ with Eye (Deprived and Undeprived) and Time (t1 and t0) as within-subjects factors and Sensory-preference in the Deprived or in the Undeprived eye as between-subjects factor revealed a significant main effect of Time (F(1,15)=13.52, p=0.002). No main effects of Eye, Sensory-Preference in the Deprived or in the Undeprived eye, nor other interaction effects were found (all ps>0.1). These findings suggest a general decrease of sensitivity at t1 (mean d’ ± SE for each session: t0 Deprived 1.44 ± 0.21; t1 Deprived 1.13 ± 0.22; t0 Undeprived 1.37 ± 0.26; t1 Undeprived 1.14 ± 0.23; see Figure 2a).

**Figure 2.**
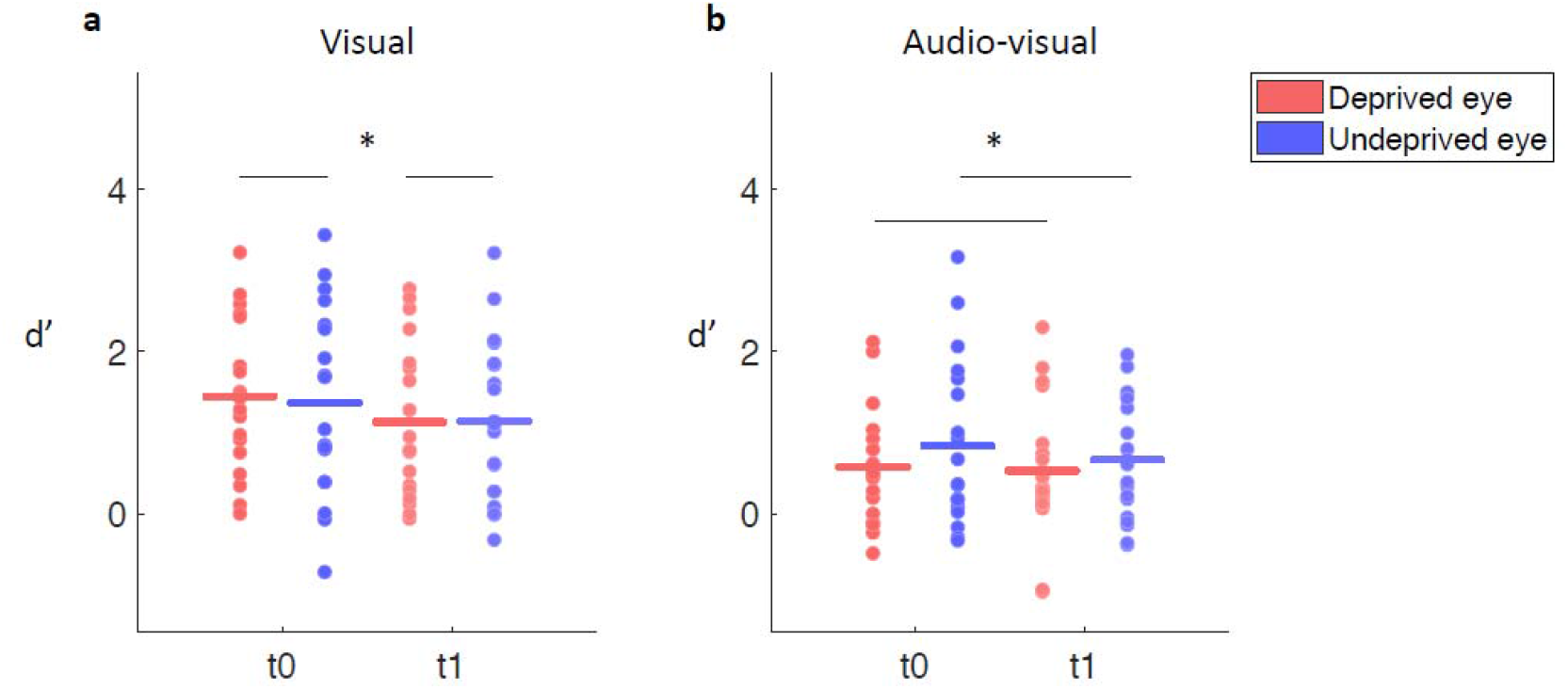
Behavioral performance computed as sensitivity (d’). (a) Both in visual and (b) audio-visual conditions, the group mean d’ is reported for t0 and t1 within each eye (Deprived and Undeprived); superimposed dots represent single subjects’ data; significant differences are highlighted with *. Sensitivity in the visual condition decreased after MD in both Deprived and Undeprived eye. Fission illusion in the audio-visual condition was greater for the Deprived than Undeprived eye.

#### Audio-visual

The mixed-design ANOVA performed on audio-visual d’ revealed a significant main effect of Eye (F(1,15)=7.53, p=0.015), showing that the Undeprived eye was less susceptible to the fission illusion (mean d’ ± SE for each session: t0 Deprived 0.58 ± 0.16; t1 Deprived 0.53 ± 0.19; t0 Undeprived 0.84 ± 0.23; t1 Undeprived 0.66 ± 0.17). A tendency towards significance emerged for the interaction between Eye, Sensory-Preference in the Deprived eye, and Sensory-Preference in the Undeprived eye (F(1,15)=4.01; p=0.064). This tendency might suggest that the participant’s Sensory-Preference had an eye-specific impact; being an *Audio*-subject or a *Visual*-subject might affect the level of the fission illusion perceived with that eye (in Deprived eye mean d’ ± SE A-group: 0.52 ms ± 0.19; V-group: 0.69 ms ± 0.20; in the Undeprived eye A-group: 0.65 ms ± 0.20; V-group: 0.96 ms ± 0.36). No significant main effects of Time, Sensory-Preference in the Deprived or in the Undeprived eye, nor other interactions emerged (all ps>0.08). Notably, a strong fission illusion was elicited in both eyes as highlighted by the small d’ measured in each of the four sessions (see Figure 2b).

Since no main effect of the Sensory-Preference emerged neither in visual nor in audio-visual ANOVAs, the analyses of EEG activity were performed on the whole group.

### Neural oscillations

To specifically investigate whether MD primarily affected feedback and/or feedforward connectivity, we assessed the impact of MD on induced and evoked neural oscillations associated with the processing of visual and audio-visual stimuli.

### Unisensory Visual

#### Induced power

The cluster-based permutation test performed on induced oscillatory activity within the low-frequency range [4-30 Hz] revealed a significant difference for *PowChangeDeprived* vs. *PowChangeUndeprived* (p<0.009) spanning from occipital to frontal regions. MD elicited a marked decrease of induced activity in the alpha range [10-16 Hz] between 0 to 0.12 seconds selectively for the Deprived eye (see Figure 3a, b). While no significant effects were found when comparing the oscillatory activity between the two eyes at t0 (all ps>0.51), the comparison performed on induced oscillatory activity measured at t1 revealed a significant difference (p<0.002) between Deprived and Undeprived eye for the power in the alpha range [10-16 Hz]. These planned comparisons confirmed a direct effect of MD, and excluded possible confounds due to differences between the eyes at baseline. To further investigate the time-course of the MD effect in the alpha range, we tested the difference between t1 and t0 for each eye (i.e., Deprived, Undeprived). For each session and participant, we extracted the mean induced alpha power [10-16 Hz] measured across three occipital electrodes (E36, E38, E40, which corresponded to the peak of the statistical effect in the *PowChangeDeprived* vs. *PowChangeUndeprived* cluster-based permutation test). A series of paired t-tests were performed between t0 and t1 for each eye, at each time-point within the whole-time window of interest [0-0.5 s] (FDR corrected, q=0.05). A significant difference was found only for the Deprived eye, showing a clear decrease in the alpha synchronization after MD (from 0 to 0.14 s; for Undeprived eye all ps>0.97; see Figure 3c).

**Figure 3.**
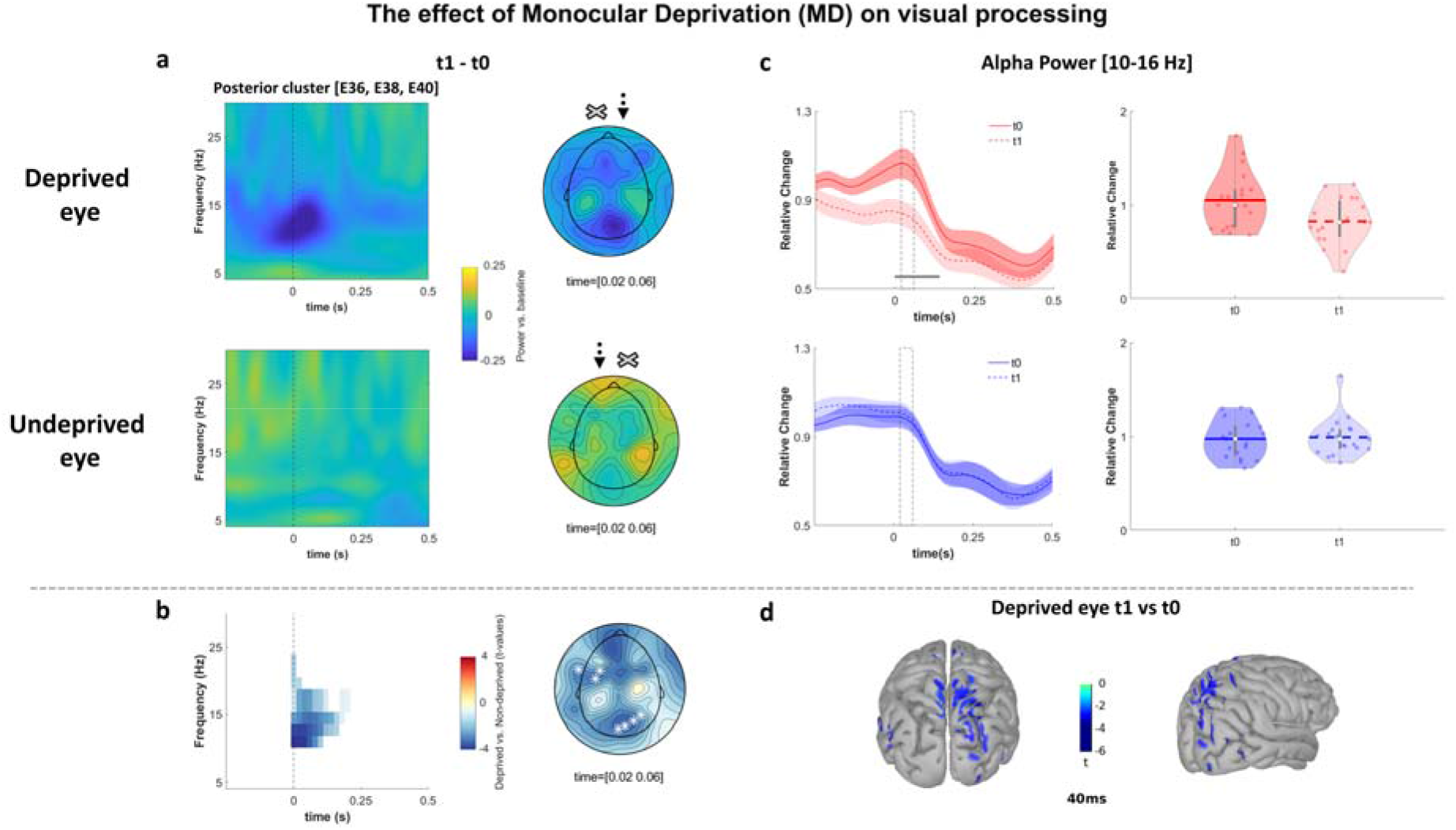
The effect of Monocular Deprivation (MD) on visual processing, visual MD effect. (a) Oscillatory activity calculated as the difference between t1 minus t0 at each eye: *PowChangeDeprived* (upper row) and *PowChangeUndeprived* (bottom row) are plotted as a function of time [−0.25 - 0.5 s] and frequency [4-30 Hz]. The plots show the average across occipital electrodes (E36, E38, E40); 0 s indicates stimulus onset. Topographies in the alpha range [10-16 Hz] at a representative time window [0.02 - 0.06 s]; arrows indicate which eye was stimulated (here depicted a participant with right-eye dominance); crosses represent the eye covered by a translucent patch. (b) Statistical results. Time-frequency plot highlighting significant differences between *PowChangeDeprived* and *PowChangeUndeprived* identified by the cluster-based permutation test (p<0.025, two-tailed) and the corresponding topography for the alpha range [10-16 Hz] at a representative time window [0.02 - 0.06 s]; electrodes belonging to the significant cluster are highlighted with white asterisks. (c) Time-course at the group level of the mean power in the alpha range [10-16 Hz] at t0 and t1 (data are averaged across electrodes E36, E38, E40) separately displayed for the Deprived and the Undeprived eye (upper and bottom rows); shaded areas represent the standard error of the mean; the continuous horizontal grey line indicates the significant difference between t0 and t1 in the Deprived eye (from 0 to 0.14 s; p<0.05, FDR corrected). The dashed grey boxes represent the time window [0.02 - 0.06 s], comprising the alpha peak, in which the power in the alpha range [10-16 Hz] was extracted for each subject (across channels E36, E38, E40) and shown in the corresponding violin plots (right side; each dot represents individual data). (d) Source analysis performed to localize the visual MD effect; the image shows, at 40 ms after stimulus onset, the area in which the power in the alpha range [10 - 16 Hz] significantly decreased at t1 with respect to t0.

No significant differences between *PowChangeDeprived* and *PowChangeUndeprived* were found in the high-frequency range [30-80 Hz] (all ps>0.67, see Figure S3).

#### Evoked power

Cluster-based permutation analyses on evoked oscillatory activity were performed contrasting *PowChangeDeprived* vs. *PowChangeUndeprived* to test whether MD alters feedforward visual processing. No significant differences emerged in either low or high frequencies (all *p*s>0.13, see Figure S4).

In sum, a visual MD effect emerged selectively for the Deprived eye and in the induced oscillatory activity within the alpha range.

### Source analysis

We investigated the electrical sources of the visual MD effect. To this end, a permutation paired t-test (1000 randomizations) was performed at the source level, on the power in the alpha range [10-16 Hz] between t0 and t1 for the Deprived eye (time window [0-0.5 s]; FDR correction was applied on the time dimension). Results revealed that the visual MD effect was mainly located within the right hemisphere and comprised the superior parietal gyrus, the superior occipital gyrus, intraparietal and subparietal sulcus, and extended to calcarine sulcus (corrected p-threshold: 0.003; see Figure 3d).

### Audio-visual

After we assessed the impact of MD on unisensory visual processing (visual MD effect), we investigated whether MD can also affect audio-visual processing at the neural level. The induced and evoked oscillatory activities were tested separately within low and high-frequency ranges.

#### Induced power

The cluster-based permutation performed on induced oscillatory activity within the low-frequency range [4-30 Hz] between *PowChangeDeprived* and *PowChangeUndeprived* showed no significant effects (all ps>0.45, see Figure S5). In contrast, the same analysis performed within the high-frequency range [30-80 Hz] revealed a significant effect in gamma activity (p<0.015). An increase in gamma activity [65-75 Hz] was found in the Undeprived eye between 0.16 and 0.26 s, mainly in posterior electrodes (see Figure 4a, b). The planned comparison showed no significant effect at t0 (all ps>0.07), while a significant difference between the two eyes emerged only at t1 (p<0.022), confirming that MD specifically guided the effect. To further investigate the time-course of the induced gamma effect, we tested the difference between t0 and t1 within each eye. For each session and participant, we extracted the mean induced gamma power [65-75 Hz] measured across two parieto-occipital electrodes (E40 and E42, which corresponded to the peak of the statistical effect in the *PowChangeDeprived* vs. *PowChangeUndeprived* cluster-based permutation test). A series of paired t-tests were performed between t0 and t1 for each eye, at each time-point within the whole-time window of interest [0-0.5 s] (FDR corrected, q=0.05). Only for the Undeprived eye, a significant effect was found (from 0.12 to 0.2 s; for Deprived eye all ps>0.5; see Figure 4c).

**Figure 4.**
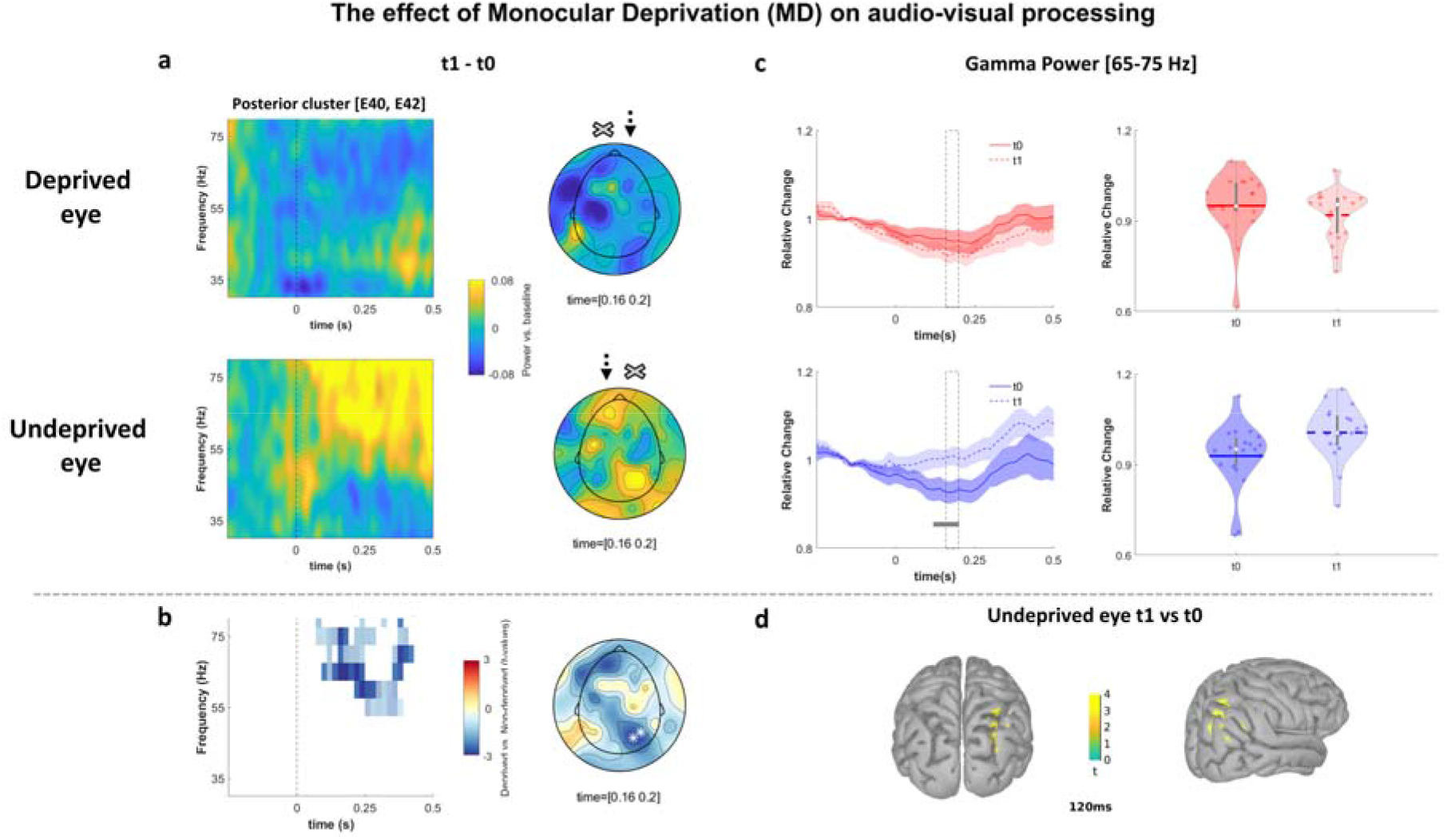
The effect of Monocular Deprivation (MD) on audio-visual processing, audio-visual MD effect. (a) Oscillatory activity calculated as the difference between t1 minus t0 at each eye: *PowChangeDeprived* (upper row) and *PowChangeUndeprived* (middle row) are plotted as a function of time [−0.25 - 0.5 s] and frequency [30-80 Hz]. The plots show the average across posterior electrodes (E40, E42); 0 s indicates the stimulus onset. Topographies in the gamma range [65-75 Hz] at representative time window [0.16 - 0.2 s]; arrows indicate which eye was stimulated (represented in a participant with a right-eye dominance), and crosses represent the eye covered by a translucent patch. (b) Statistical results. Time-frequency plot highlighting significant differences between *PowChangeDeprived* and *PowChangeUndeprived* identified by the cluster-based permutation test (p<0.025, two-tailed) and corresponding topography for the gamma range [65-75 Hz] at a representative time window [0.16 - 0.2 s]; electrodes belonging to the significant cluster are highlighted with white asterisks. (c) Time-course at the group level of the mean power in gamma range [65-75 Hz] at t0 and t1 (data are averaged across electrodes E40, E42) separately displayed for the Deprived and the Undeprived eye (upper and bottom rows); shaded areas represent the standard error of the mean; the continuous horizontal grey line indicates a significant difference between t0 and t1 for the Undeprived eye (from 0.12 to 0.2 s after stimulus onset; p<0.05, FDR corrected). The dashed grey boxes represent the time window [0.16 - 0.2 s] in which the power in the gamma range [65-75 Hz] was extracted for each subject (across channels E40 and E42) and shown in the corresponding violin plots (right side). (d) Source analysis performed to localize the audio-visual MD effect; the image shows at 120 ms after stimulus onset the area in which the power in the gamma range [65-75 Hz] significantly increases at t1 with respect to t0.

The gamma band increase during audio-visual processing in the Undeprived eye following MD might suggest an upweighting of the auditory modality. If this would be the case, a greater neural response in the gamma range to unimodal acoustic stimulation (A and AA condition) should emerge selectively for the Undeprived eye after MD. To this end, a hypothesis-driven (p<0.05, one tail) cluster-based permutation test was performed on the induced oscillatory activity in response to auditory stimuli (average across A and AA trials) in the high-frequency range [30-80 Hz] across all electrodes, frequencies, and time-points [0-0.5 s], between t0 and t1, separately in each eye. The analysis revealed a significant increase in gamma activity between 100 and 300 ms in response to auditory stimulation after MD (p<0.04), selectively for the Undeprived eye (see Figure 5). Conversely, no significant difference emerged for the Deprived eye (p>0.35). This significant effect emerged in parietal electrodes as for the audio-visual MD effect. These findings support our hypothesis that the increased induced gamma activity during audio-visual processing is due to an upweighting of auditory input.

**Figure 5.**
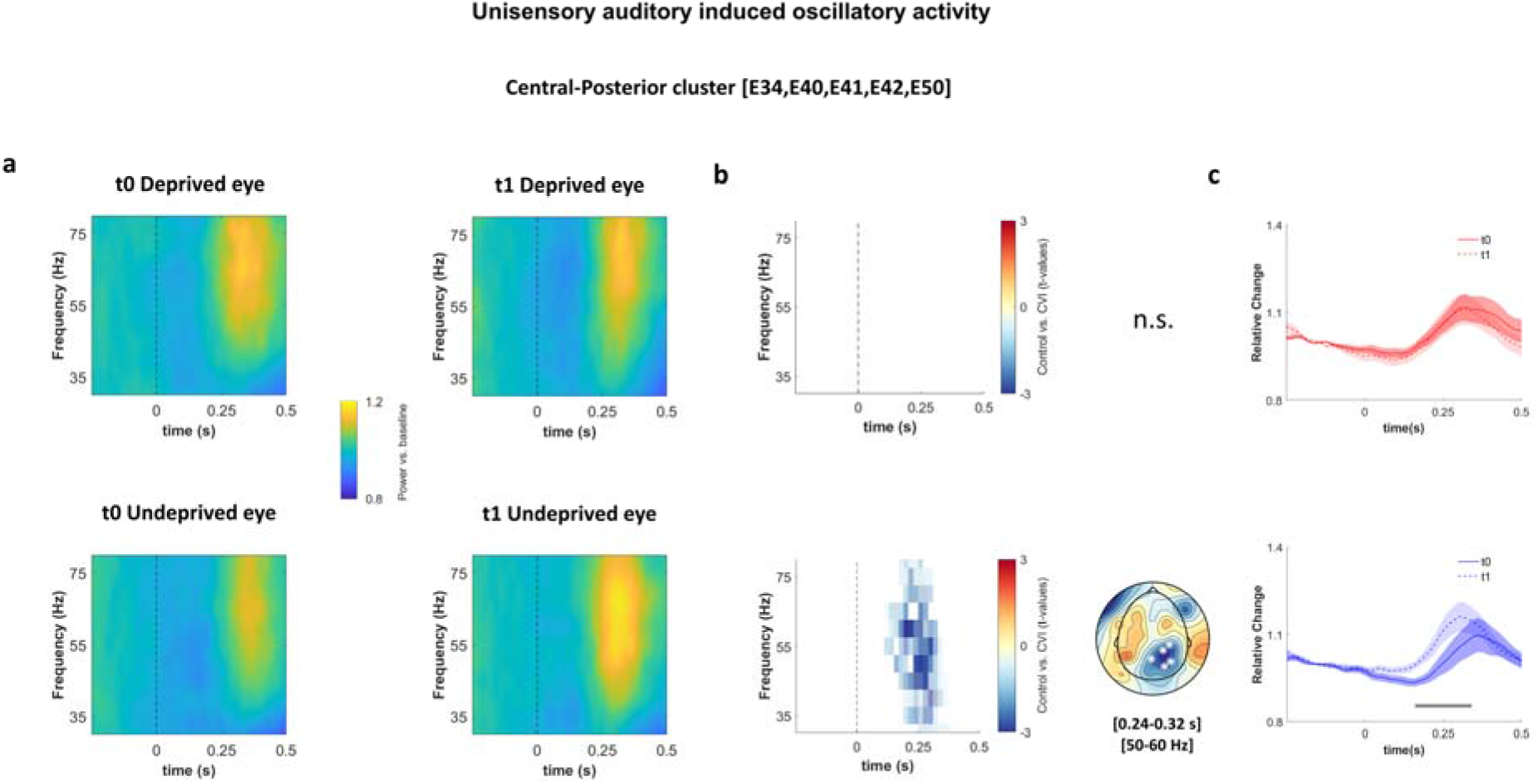
Increased induced gamma activity after MD during unisensory auditory processing. (a) Oscillatory activity at t0 and t1 are plotted as a function of time [−0.25 - 0.5 s] and frequency [30-80 Hz] for the Deprived (upper row) and Underprived eye (bottom row). The plots show the average across central-posterior electrodes (E34, E40, E41, E42, E50); 0 s indicates the stimulus onset. (b) Results of the cluster-based permutation test are shown in the time-frequency plots and highlight significant differences between t0 and t1, which emerged only for the Undeprived eye. Topography shows the results of the cluster-based permutation test for the Undeprived eye in the gamma range [50-60 Hz] over a representative time window [0.24 - 0.32 s]; white asterisks highlight significant electrodes (E34, E40, E41, E42, E50). (c) Time-course at the group level of the mean power in gamma range [50-60 Hz] at t0 and t1 (data are averaged across electrodes E34, E40, E41, E42, E50), separately displayed for the Deprived and the Undeprived eye (upper and bottom rows); shaded areas represent the standard error of the mean; the continuous horizontal grey line indicates a significant difference between t0 and t1 for the Undeprived eye (from 0.16 to 0.34 s after stimulus onset; p<0.05, FDR corrected).

#### Evoked power

When the difference between *PowChangeDeprived* and *PowChangeUndeprived* was tested in the evoked power, no significant difference was found within the low-frequency range (all ps>0.09), nor within the high-frequency range (p>0.05, see Figure S7).

To sum up, the audio-visual MD effect selectively emerged for the Undeprived eye in the induced gamma power. This neurophysiological change seems to be driven by increased responsiveness to auditory inputs.

### Source analysis

We investigated the sources of the audio-visual MD effect. To this end, a permutation paired t-test (1000 randomizations) was performed at the source level, on the power in the gamma range [65-75 Hz] between t0 and t1 for the Undeprived eye (time window [0-0.5 s]; FDR correction was applied on the time dimension).

Results revealed that audio-visual MD effect was mainly located at the cortical level around the right intraparietal sulcus (corrected p-threshold: 0.002; see Figure 4d).

### Association between neural and behavioral changes due to MD

For visual and audio-visual conditions, we investigated the degree of association between brain activity alterations and changes in behavioral performance. The average power at the frequencies of interest was extracted within a time window and across channels that resulted significant in the cluster-based permutation tests (visual and audio-visual MD effects). At the individual level, we computed the normalized difference between t1 and t0 within each eye (*[PowChangeDeprived/t0 Deprived] * 100; [PowChangeUndeprived/t0 Undeprived] * 100*). Shapiro-Wilk tests confirmed that data in each condition were normally distributed (all ps >0.05). Thus, within each condition (i.e., visual and audio-visual) and for each eye (i.e., Deprived and Undeprived), the power change was correlated with the corresponding change in sensitivity between t0 and t1 using Pearson’s correlation coefficient.

### Unisensory Visual

We first tested whether observed changes of oscillatory activity in induced alpha power following MD (visual MD effect) were associated with changes in behavioral performance. To this aim, we extracted the average power in the 10-16 Hz range between 0.02 and 0.06 s from the three significant electrodes (E36, E38, and E40) for each session. Then, the normalized t1-t0 difference of *PowChangeDeprived* and *PowChangeUndeprived* was computed.

A significant positive correlation between normalized *PowChangeDeprived* and visual sensitivity change in the Deprived eye (d’ change) was found (r(18) = 0.492, p=0.038). Following MD, the visual sensitivity in the Deprived eye decreased in parallel with a reduction in induced alpha power in the same eye. When the correlation between induced alpha change and visual sensitivity change was tested for the Undeprived eye, no significant effect was found (p>0.99).

### Audio-visual

Next, we tested whether the increase in induced gamma after MD period (audio-visual MD effect) was associated with illusory fission percept. Thus, within each session, the induced power between 65 and 75 Hz was extracted within the 0.16-0.2 s time window and across the two significant posterior channels (E40 and E42). Then, the normalized t1-t0 difference of *PowChangeDeprived* and of *PowChangeUndeprived* was computed. No significant correlation was found neither for the Deprived eye nor for the Undeprived eye (all ps>0.43). However, at the behavioral level, a tendency for aninteraction between illusion perception and sensory preferences was found (F(1,15)=4.01, p=0.06), indicating that *Visual* subjects tended to experience less illusion than *Audio* subjects, especially in the Undeprived eye (see section Audio-visual in the Behavioral Results). Interestingly, the perception of the multisensory input is known to partially depend on individual sensory predisposition (e.g., Giard and Perronet, 1999; Hong et al., 2021). Therefore, despite this was not the focus of the paper, we additionally explored whether the association between neural and behavioral changes might be affected by individual sensory preference. Specifically, we calculated the correlation between *PowChangeUndeprived* and audio-visual d’ change in the Undeprived eye within *Audio* and *Visual* groups, classified according to participants’ Sensory-Preference (estimated for the same eye). Pearson correlation revealed a significant negative correlation in the V-group (r(5) = −0.937; p=0.019), while a tendency toward a significant positive correlation was found in the A-group (r(13) = 0.513; p=0.073). The two correlations significantly differed (difference=1.45 CI [0.17 1.80]. Since Pearson correlations were performed, the confidence interval was adjusted as described in Wilcox (2009); see Supplementary Materials Figure S8). While these results are based on an explorative analysis and small sub-samples, they seem to suggest that the audio-visual MD effect might have a different impact on the illusory percept according to the participants’ Sensory-Preference: in V-group, the increase of induced gamma activity in the Undeprived eye was positively associated with a fission illusion increase (smaller d’ after MD), while in A-group the increase of induced gamma activity seemed to be associated with a fission illusion decrease (larger d’ after MD).

## Discussion

In this study, adult neural plasticity of both unisensory visual and multisensory audio-visual processes was investigated to assess whether the impact of MD extends beyond visual processing. Induced and evoked oscillatory activity changes both in visual and audio-visual processing were measured after 150 minutes of altered visual experience (brief MD). Induced alpha associated with the early phase of visual processing (<150ms) decreased after MD selectively for the Deprived eye. Conversely, induced gamma associated with audio-visual processing increased after MD only in the Undeprived eye, within a later temporal window (~100-300ms). Notably, both visual and audio-visual processing alterations were found selectively for the induced component of neural oscillations (see Figure 6). Source modeling linked both the visual and audio-visual MD effects to the right parieto-occipital cortex. Our data reveal the specific neural signatures of temporary MD effects on visual and audio-visual processes and shed light on their shared feedback nature. We demonstrated that a brief period of monocular visual experience in adulthood specifically changes the neural response to multisensory audio-visual events because of plasticity in feedback connectivity.

**Figure 6.**
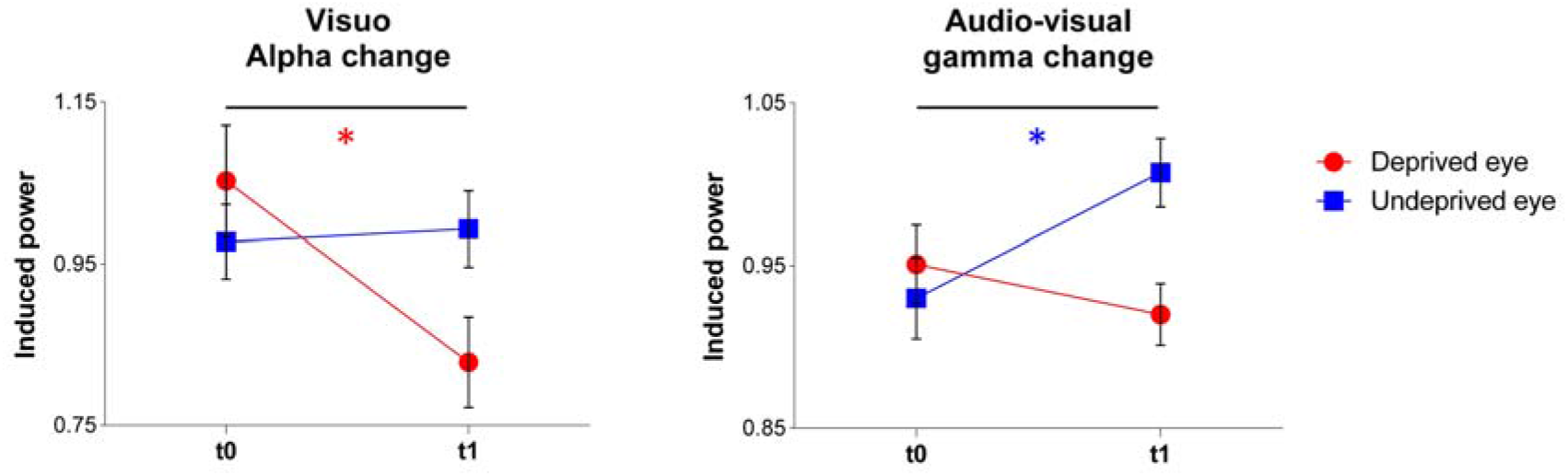
Visual and audio-visual MD effects on induced oscillatory activity. The plots show the % normalized power change due to MD in unisensory visual and multisensory audio-visual processing. On the left, the visual MD effect: decreased induced alpha power selectively for the Deprived eye (red) during unisensory visual processing. On the right, the audio-visual MD effect: enhanced induced gamma power selectively for the Undeprived eye (blu) during audio-visual processing. Each dot represents the group mean and bars the SE. Significant differences are highlighted with * (red indicates effect for the Deprived eye and blu for the Undeprived eye).

### Spectro-temporal properties of the visual MD effect

The observed visual MD effect is in line with previous studies (Lunghi et al., 2015a, 2015b; Zhou et al., 2015; Binda et al., 2018; Schwenk et al., 2020). The reduced alpha synchronization observed when the task was performed with the Deprived eye after MD is consistent with increased excitability to compensate for the absence of stimulation during the deprivation phase. Source modeling suggested that the alpha reduction was mainly localized in the right hemisphere in superior parieto-occipital areas, with some activities extending to primary visual areas (calcarine sulcus). These results confirm the modulation of alpha rhythm induced by short-term plasticity, previously shown in the frequency domain as a change in alpha peak amplitude (Lunghi et al., 2015a), and characterize the spectro-temporal properties of this neural effect. Namely, a selective decrease of alpha synchronization [10-16 Hz] occurs in the early stages of visual processing (<150ms). Notably, only induced activity (i.e., not phase-locked to stimulus onset) was altered by MD, revealing the feedback nature of visual short-term plasticity. Overall, these results are coherent with spectroscopy data showing an increase of excitability in the early visual cortex as indicated by reductions of inhibitory gamma-aminobutyric acid (GABA) concentration after short-term MD (Lunghi et al., 2015b). Moreover, selectively for the Deprived eye, a significant correlation was found between the decrease in induced alpha activity and the decrease in visual sensitivity after MD, suggesting a potential link between this neural change and the ability to discriminate temporal aspects of visual processing.

### The impact of MD on audio-visual processing

Previous behavioral studies have shown that multisensory perception can be altered by MD (Lo Verde et al., 2017; Opoku-Baah and Wallace, 2020). By measuring changes in neural oscillations, we assessed neural mechanisms underpinning multisensory short-term plasticity and revealed the specific enhancement of induced gamma activity [65-75 Hz] selectively for the Undeprived eye when processing audio-visual input. The audio-visual MD effect involved the right intraparietal sulcus, suggesting its central role in short-term plasticity induced by MD.

We hypothesized that the audio-visual MD effect could be due to a crossmodal upweighting of the other modality (i.e., audition). To this end, we tested whether the neural response to unimodal auditory stimuli was increased following MD in each eye. The observation that gamma response to auditory input increased following MD selectively for the Underprived eye confirmed our hypothesis. This result revealed that short-term plasticity following MD alters both visual and auditory neural representations. Interestingly, alterations of neural excitability due to temporary binocular deprivation were previously shown to increase heteromodal responses in the visual cortex (Merabet et al., 2008).

From a neurophysiological perspective, it is important to remark that neural profiles of short-term plasticity following MD seem to depend on the type of input at hand. While the visual stimulus was the same in visual and audio-visual conditions, short-term plasticity was characterized by specific oscillatory fingerprints (Siegel et al., 2012): alpha decreased during visual processing while gamma increased during audio-visual and auditory processing.

To what extent does this neural alteration interact with individuals’ sensory predisposition? In the Undeprived eye, the correlation between enhanced gamma activity in audio-visual processing and behavioral performance seems to indicate opposite MD impacts on illusory perception with respect to participants’ Sensory-Preference. Although preliminary, this result opens the possibility that the upweighting of auditory information during audio-visual processing after MD affects visual perception according to individual sensory preference. Coherently with the extreme flexibility and adaptability of multisensory functions (Giard and Perronet, 1999; Fujisaki et al., 2004; Van Atteveldt et al., 2014), the perception of the multisensory input after MD might change as a function of individual sensory predisposition. This inference should be further verified with psychophysics experiments (see Rohe et al., 2019), designed to directly estimate auditory modality’s weight changes during audio-visual processing after MD. Interestingly, a recent study investigating cross-modal recalibration highlighted how individual variability in one sensory modality (visual reliability) differently affects recalibration of the other modality (audition) (Hong et al., 2021).

Binocular input was shown to be critical for developing audio-visual perception: anomalies in different audio-visual perceptual tasks were reported in cases of monocular enucleation (Moro and Steeves, 2018a, 2018b), individuals affected by early monocular cataracts (Chen et al., 2017), and people suffering from amblyopia (Narinesingh et al., 2015, 2017; Richards et al., 2017). The present results, revealing that MD induces short-term plasticity of audio-visual processing, encourage possible treatments of audio-visual anomalies associated with MD. Increasing evidence in animal studies supports clinical treatment of adult amblyopia (Hensch and Quinlan, 2018), and crucially a recent study conducted with adult people affected by amblyopia has shown that MD combined with physical exercise could be a promising clinical intervention to promote visual recovery (Lunghi et al., 2019). Future studies might help understanding whether multisensory audio-visual processing could also benefit from this novel clinical treatment.

### The pivotal role of induced cortical response in experience-dependent plasticity

Both visual and audio-visual MD effects were selectively found in induced neural oscillations, likely reflecting main alterations in feedback processing integrating sensory input and ongoing cortical activity (Galambos 1992; Klimesch et al., 1998; Tallon-Baudry and Bertrand, 1999; Chen et al., 2012; Keil et al., 2022). Instead, the evoked phase-locked activity (feedforward), related to the processing within the thalamo-cortical pathway (Galambos 1992; Keil et al., 2022), was not affected.

Visual MD effect was found in the induced alpha band, which was previously demonstrated to be drastically impaired by the transient absence of visual experience during development (Bottari et al., 2016). Animal studies involving congenital deaf cats demonstrated that induced and not evoked oscillatory activity in the auditory cortex is extremely reduced across a wide range of frequency bands (e.g., Yusuf et al., 2017). The authors hypothesized that the absence of sensory stimulation prevents the development of neural mechanisms allowing the integration of sensory signals and internal representations. Evidence of alterations of induced oscillatory activity following sensory deprivation was reported not only within modality, for visual (Bottari et al., 2016) and auditory systems (Yusuf et al., 2017), but also cross-modally. In humans, early-onset deafness selectively affects induced oscillatory activity associated with visual processing (Bednaya et al., 2021). The present audio-visual MD effect (increased induced gamma activity) provides evidence in the same direction also for multisensory processing. Taken together, this evidence suggests a substantial alteration of cortico-cortical feedback activities, in case of sensory input absence in both developmental and adult brain, for unisensory and multisensory functions. This is in line with previous evidence suggesting that the plasticity of feedback connectivity represents an extremely flexible mechanism to process sensory information according to changing demands (Polley et al., 2006).

### Limitations of the study

Here, we focused on neural oscillations changes, and given the time constraints intrinsic in the MD effect (which is maximum within the first 15 minutes following the deprivation, Lunghi et al., 2011), we could not insert a direct measure of ocular dominance, which is usually assessed with binocular rivalry (e.g., Lunghi et al., 2011). Thus, we relied on eye-specific oscillatory activity alterations selectively at t1 (and their lack at t0) as indications of the change in interocular excitability balance: the decreased alpha during visual processing is compatible with the strengthening of the Deprived eye after MD. While a clear reduction of strength for the Undeprived eye (i.e., increased alpha activity) did not emerge, this is in line with the literature: the strongest impact of MD is known to be on the Deprived eye, while the opposite effect on the Undeprived eye was found to be much smaller (Lunghi et al., 2015a; Binda et al., 2018) or even absent (Zhou et al., 2015). Thus, it is possible that larger sample sizes are required to measure neural effects on the Undeprived eye during visual processing. Further studies might help to assess whether depriving the non-dominant eye will lead to the same MD effects. However, given the absence of difference at t0, we can rule out the possibility that our effects are due to baseline differences between the dominant and the non-dominant eye.

Noticeably, the visual MD effect emerged here for the Deprived eye, while the audio-visual MD effect emerged only for the Undeprived eye. While these effects are in line with previous behavioral studies (Lo Verde et al., 2017; Opoku-Baah and Wallace, 2020), they also support a crucial role of MD in inducing flexible alterations of interocular excitability balance and, in turn, in audio-visual processes. Moreover, although an interaction between MD and eye dominance cannot be excluded, a recent study reported that the MD effect was the same regardless of whether the dominant or the non-dominant eye was deprived (Schwenk et al., 2020).

Finally, while the interaction between sensory preference and gamma change in audio-visual processing seems of interest, a larger sample with a balanced number of subjects with visual and auditory sensory preferences is needed to confirm our intriguing preliminary result on how multisensory plasticity can be affected by individual sensory predisposition.

## Conclusions

These results demonstrated that a brief period of monocular visual experience in adulthood is able to change the neural response not only to visual stimuli but also to multisensory events. The data unveiled the spectral fingerprints of adult short-term plasticity induced by a brief period of MD for visual and audio-visual processing. We found enhanced excitability (i.e., decreased induced alpha activity) for the Deprived eye during an early phase of visual processing, and we demonstrated the presence of neural alterations beyond the visual processing. Gamma activity associated with audio-visual processing was increased by MD at a later latency and only when the task was performed with the Undeprived eye. The analyses of responses to unimodal auditory input indicate an upweighting of sound input following MD, selectively for the Undeprived eye. Importantly, these distinct neural effects were selectively found in the induced neural oscillations, revealing that experience-dependent plasticity mainly involves alterations in feedback processing not only during development but also in adulthood. This observation is consistent with the existence of a general mechanism shared across sensory modalities and across the life cycle.

## Acknowledgments

This work was funded by PRIN 2017 research grant (Prot. 20177894ZH) to Davide Bottari.

We would like to thank Cesare V. Parise for the helpful discussion and feedback regarding data analysis.

## CRediT authorship contribution statement

Alessandra Federici: Conceptualization, Investigation, Data curation, Formal analysis, Writing - original draft, Writing - review & editing. Giulio Bernardi: Conceptualization, Writing - review & editing. Irene Senna: Formal analysis, Writing - review & editing. Marta Fantoni: Investigation. Marc Ernst: Formal analysis, Writing - review & editing. Emiliano Ricciardi: Resources, Writing - review & editing. Davide Bottari: Conceptualization, Investigation, Data curation, Formal analysis, Writing - original draft, Writing - review & editing, Supervision, Project administration, Funding acquisition.

