## Supplementary Materials for "Short-term monocular deprivation boosts neural responsiveness to audio-visual events for the undeprived eye"

**Simple size estimation for the main EEG experiment**

We estimated the minimum sample size needed to replicate previous evidence of neural MD effect on visual processing. Starting from the previous electrophysiological (EEG) studies (Lunghi et al., 2015a), we expected the effect to be in the alpha range [8-14 Hz], in the first stages of the visual processing [0-120 ms], and in occipito-parietal electrodes (E33, E34, E36, E38). As planned for our main analyses, we contrasted the difference between t1 and t0 in the Deprived and Underpived eye. Using the data of four pilot subjects, through simulations suited for cluster-based permutation tests (500 randomization; [Wang and Zhang, 2021](https://doi.org/10.1111/psyp.13775)), we estimated a minimum sample size of 17 subjects to reach a power of 0.8 (lower threshold). Here, the final sample comprised 19 behavioral and 19 EEG datasets.

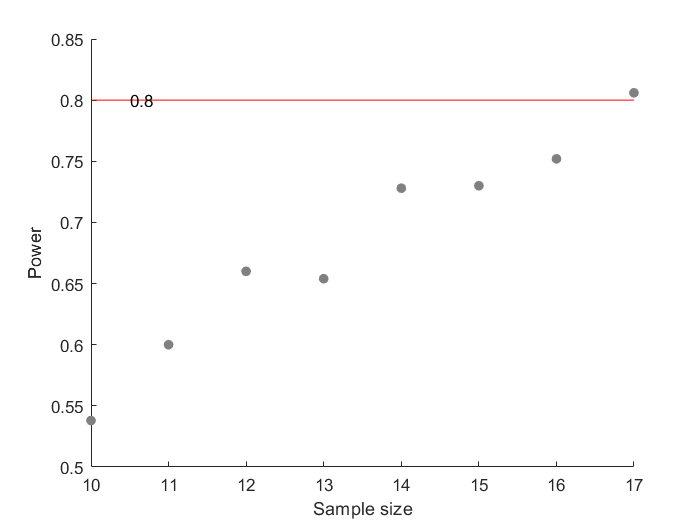

**Figure S1. Sample Size Estimation.** Expected statistical power at each sample size, starting from N=10, computed through simulations of time-frequency data on the total power (alpha = 0.05). Results showed that to achieve a power of 0.8 (red line), a minimum sample size of N=17 was required.

**Participants excluded after the preliminary behavioral assessment**

Five participants were excluded as they did not meet the inclusion criteria of the study: one perceived the fission illusion only in a few trials (20% illusory rate), and the others were entirely biased by the sound on their visual perception (>95% illusory rate). Moreover, one participant was excluded because they could not comply with instructions (difficulties in responding within the response time window).

**Conditions trials**

In fission illusion (AVA condition), the visual stimulus was presented between the two auditory stimuli, with the inter-stimulus-interval (ISI) between each stimulus being a single frame (i.e., 17 ms). The same was done for fusion illusion (VAV condition): the audio stimulus was presented within the two visual stimuli (ISIs between the stimuli: 17 ms). Instead, in coherent multisensory conditions (i.e., AV and AVAV), each audio-visual couple was presented simultaneously. In the AVAV condition, the two couples of stimuli were presented with an ISI of 51 ms (that corresponds to three frames, 17 ms × 3), as in the conditions with two unisensory stimuli (i.e., VV, AA).

**Porta Test**

To assess participant’s eye dominance, we used the Porta test. With both eyes open, the participant extends one arm and aligns the thumb finger with a distant object. Then participant alternates closing the eyes, when the thumb appears to “move” it means that the closed eye is the dominant one (<https://web.archive.org/web/20080215220943/http://www.sportvue.com/support/dominance.php>).

**Staircase procedure on the AVA condition**

The contrast level of the grey dot presented on the screen was selected for each individual via a one-up one-down staircase procedure. In the staircase procedure, 133 contrasts for the grey dots were implemented (linearly spaced). When the subject’s response corresponded to the perception of the fission illusion (i.e., response “two flashes”), the contrast decreased, while it increased after a non-illusion perception (i.e., response “one fash”). Wrong answers (i.e., response zero flashes) were not taken into account by the staircase procedure, and the same contrast was displayed again in the subsequent AVA trial. AVA condition trials were presented randomly with all the other conditions (i.e., A, AA, V, VV, VAV, AV, AVAV; maximum 15 trials for each condition). No more than three consecutive AVA trials could be displayed for a maximum of 99 trials. Flash contrast was randomly selected between 10 pre-selected contrasts in all the other conditions. The staircase procedure was considered completed when 16 reverses, changes of direction in the staircase, were reached. The threshold contrast value was computed as the mean of the last three levels of contrast that caused a reverse.

**Speeded object recognition task**

The speeded object recognition task was performed monocularly with the dominant (Deprived) eye and the non-dominant (Undeprived) eye in a randomized order.

We created two different objects: one was characterized by a small circle (1.7° diameter grey dot) and a high pitch sound (quadratic beep with a 3.8 kHz frequency and a sampling rate of 44.1 kHz). While the other by a bigger circle (2° diameter grey dot) and a lower pitch sound (quadratic beep with a 3.5 kHz frequency and a sampling rate of 44.1 kHz). The visual and auditory features were coupled in accordance with previously reported crossmodal correspondences between auditory pitch and size of visual objects (see review Spence, 2011). In each trial, participants were presented with one of the two objects defined either by visual feature alone (unisensory visual condition), auditory feature alone (unisensory auditory condition), or by the combination of auditory and visual features, and they were asked to recognize as soon as possible which of the two objects was presented. Twenty trials for each condition (only audio, only video, audio-visuo × 2 objects) were randomly presented. Computing the median reaction times (RTs) at the single-subject level, only on correct trials and excluding the outliers above or below 3 SD from the median of each condition, we compared RTs in unisensory visual and unisensory auditory trials to define the Sensory-Preference of each participant. Subjects with shorter RTs in unisensory visual trials were considered *Visual*, while those who were faster in unisensory auditory trials were considered *Audio*.

**Behavioral results in all conditions**

We reported here the mean of responses in each condition for each session. In all four sessions, we found an expected performance: when zero stimuli were presented, the means were close to zero, and the mean response increased when the number of the visual stimuli increased (maximal when 2 Visual and 2 Auditory stimuli were presented); participants are clearly able to perform the task in all sessions. Notably, the fission illusion was elicited in all sessions; indeed, AVA cell (1 Visual and 2 Auditory stimuli) is always whiter than AV cell (1 Visual and 1 Auditory stimulus).

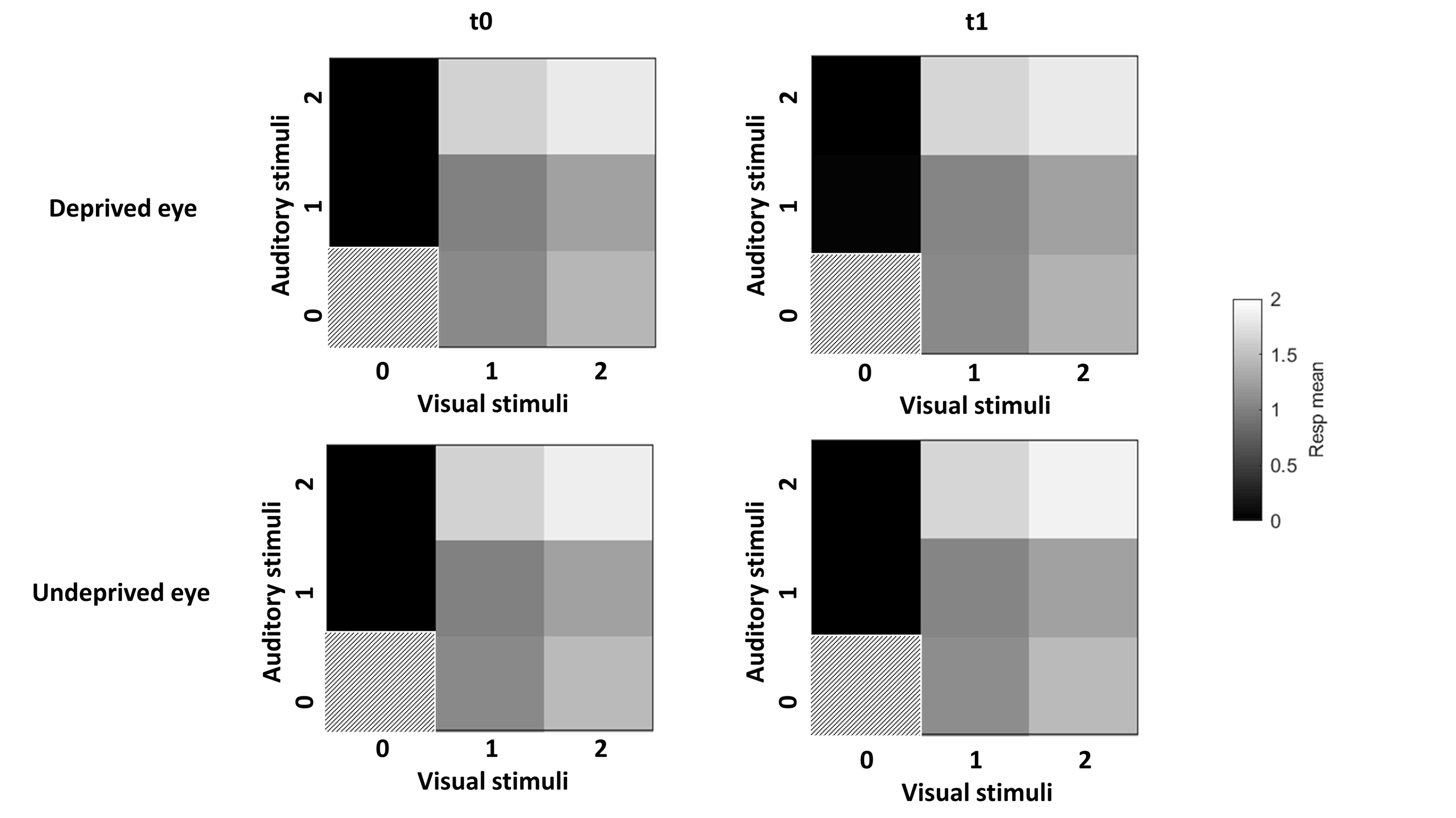

**Figure S2**. **Mean of response for each condition in each session.** The heatmaps of the four sessions (t0 Deprived, t0 Undeprived, t1 Deprived, t1 Undeprived) are shown at the group level. Each square represents one of the eight possible conditions, according to the stimulations delivered (the condition zero-zero is absent in our experiment, for this reason, we fill this square up with dashed lines). The color code represents the mean responses: black = zero is the minimum, and white = two is the maximum.

**Unisensory visual**

***Induced power*.** No significant differences between *PowChangeDeprived* and *PowChangeUndeprived* were found in the high-frequency range (all ps>0.6).
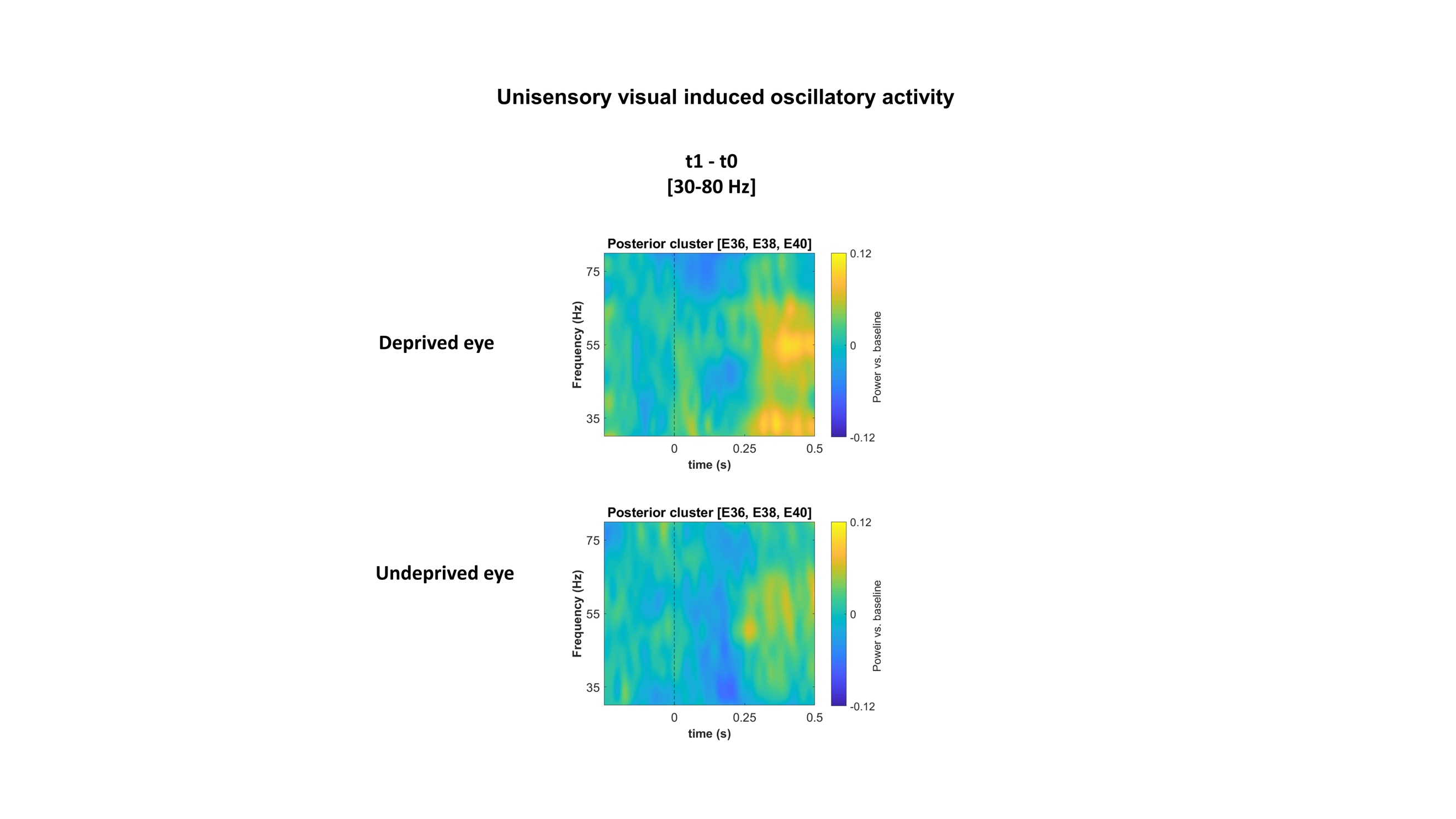
**Figure S3**. **Time-frequency plots showing high-frequency induced oscillatory activity during visual processing.** Oscillatory activity, calculated as the difference between t1 and t0 at each eye (upper row, *PowChangeDeprived*, t1 minus t0 at the Deprived eye, and bottom row, *PowChangeUndeprived*, t1 minus t0 at the Undeprived eye), are plotted as a function of time [-0.25 - 0.5 s] and frequency [30-80 Hz]. The data represent the average across occipital electrodes (E36, E38, E40); the dashed line at 0 s indicates the stimulus onset.

***Evoked power*.** No significant differences emerged in either low or high frequencies (all *p*s>0.13).

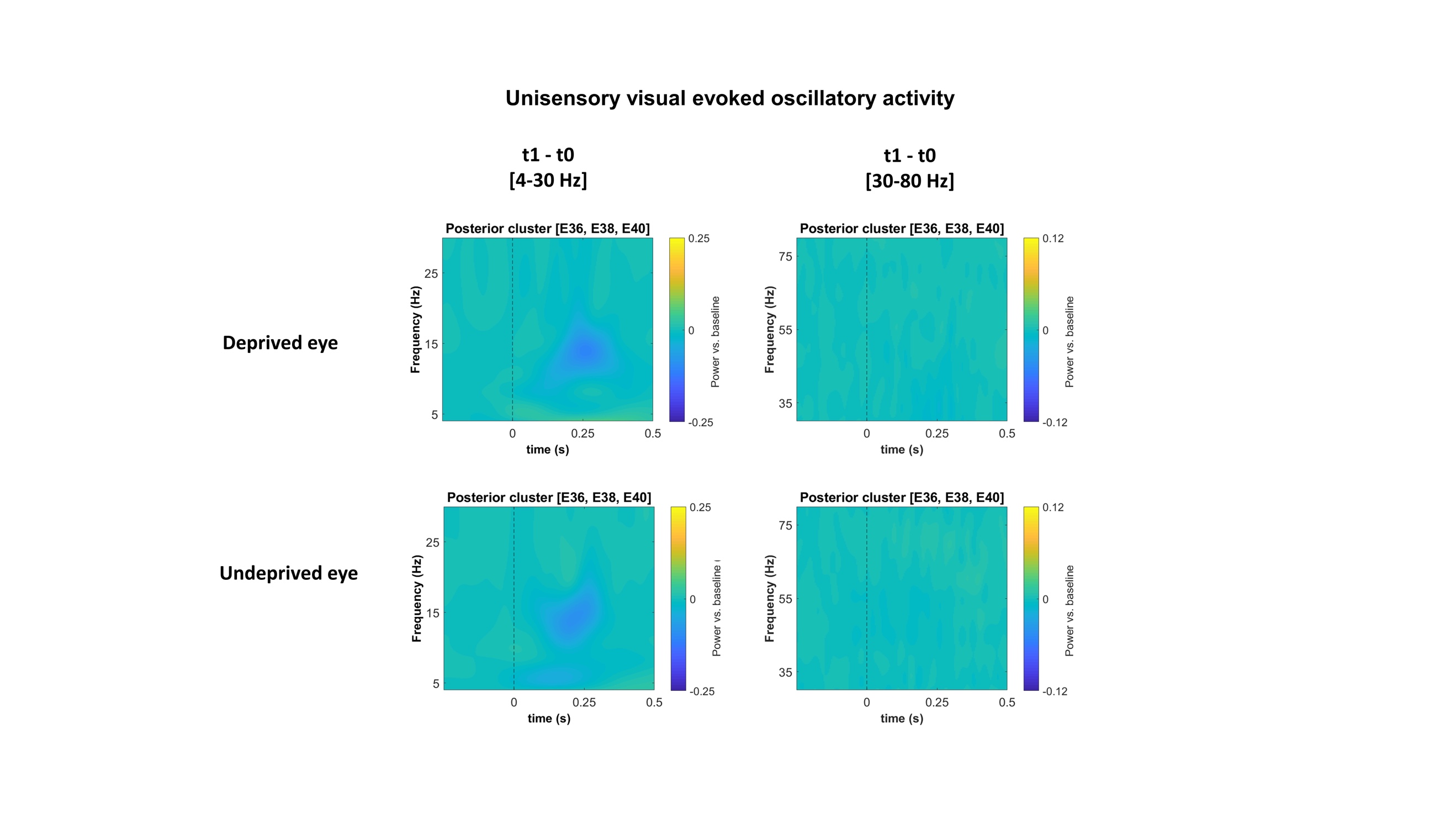
 **Figure S4. Time-frequency plots showing evoked oscillatory activity during visual processing.** Oscillatory activity, calculated as the difference between t1 and t0 at each eye (upper row, *PowChangeDeprived*, t1 minus t0 at the Deprived eye and bottom row, *PowChangeUndeprived*, t1 minus t0 at the Undeprived eye), are plotted as a function of time [-0.25 - 0.5 s] and frequency [4-30 Hz] and [30-80 Hz] (left and right column, respectively). The data represent the average across occipital electrodes (E36, E38, E40); the dashed line at 0 s indicates the stimulus onset.

**Audio-visual**

***Induced power*.** The cluster-based permutation performed between *PowChangeDeprived* and *PowChangeUndeprived* within the low-frequency range showed no significant effects (all *ps*>0.45).

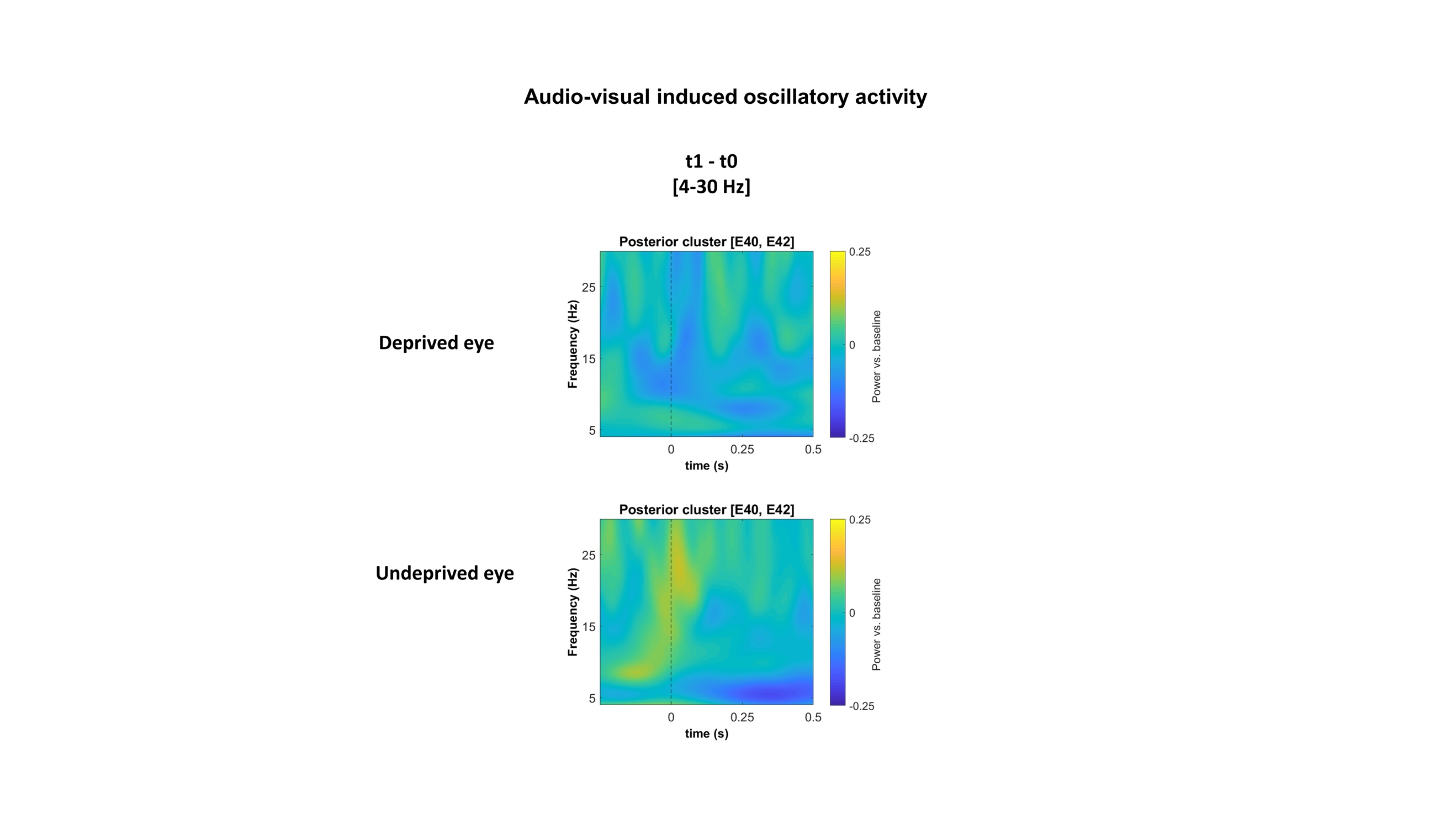
 **Figure S5**. **Time-frequency plots showing low-frequency induced oscillatory activity during audio-visual processing.** Oscillatory activity, calculated as the difference between t1 and t0 at each eye (upper row, *PowChangeDeprived*, t1 minus t0 at the Deprived eye and bottom row, *PowChangeUndeprived*, t1 minus t0 at the Undeprived eye), are plotted as a function of time [-0.25 - 0.5 s] and frequency [4-30 Hz]. The data represent the average across two posterior electrodes (E40, E42); the dashed line at 0 s indicates the stimulus onset.

***Evoked power*.** When we tested the difference between *PowChangeDeprived* and *PowChangeUndeprived* in the evoked power, no significant difference was found within the low-frequency range (all ps>0.09) nor within the high-frequency range (p>0.05).

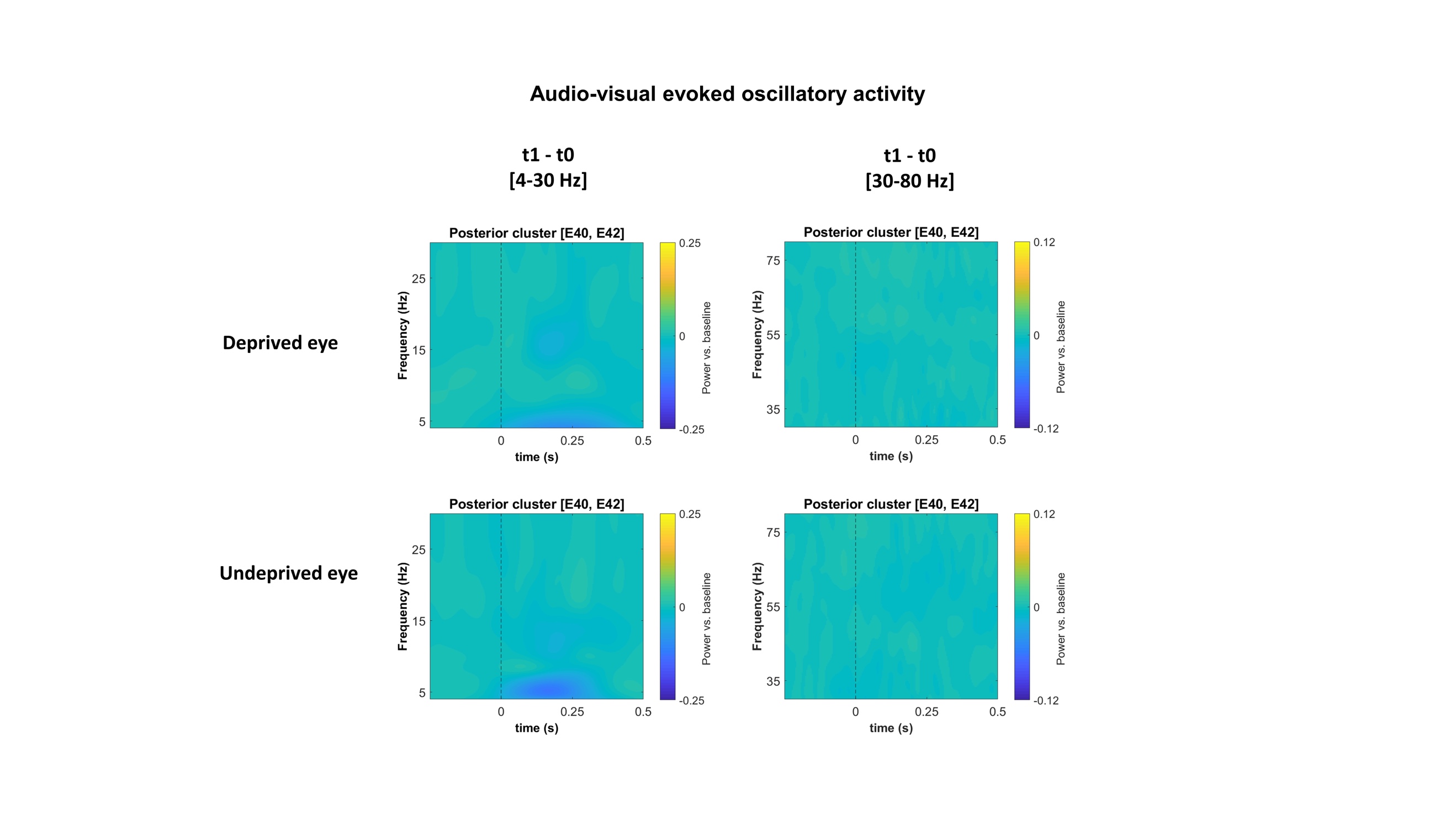

**Figure S7**. **Time-frequency plots showing evoked oscillatory activity during audio-visual processing.** Oscillatory activity, calculated as the difference between t1 and t0 at each eye (upper row, *PowChangeDeprived*, t1 minus t0 at the Deprived eye and bottom row, *PowChangeUndeprived*, t1 minus t0 at the Undeprived eye), are plotted as a function of time [-0.25 - 0.5 s] and frequency [4-30 Hz] and [30-80 Hz] (left and right column, respectively). The data represent the average across two posterior electrodes (E40, E42); the dashed line at 0 s indicates the stimulus onset.

**Comparison between correlations in the audio-visual condition**

After computing for A- and V- groups the correlations between normalized *PowChangeUndeprived* and d’ change for the audio-visual condition in the Undeprived eye, we tested by means of the bootstrap method (Pernet, Wilcox & Rousselet, 2012) whether the difference between the two correlations was significantly different from zero. Since Pearson correlations were performed, the confidence interval was adjusted as described in Wilcox (2009).

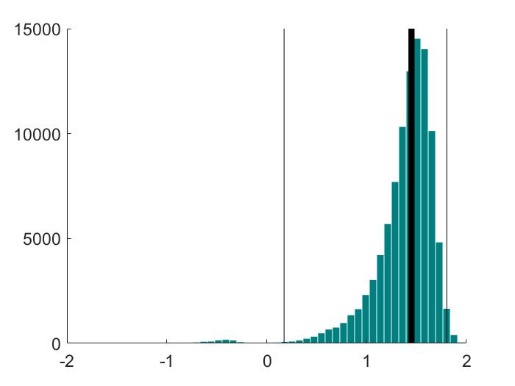

**Figure S8**. **Difference between correlations in the *Audio* and *Visual* groups**. The graph shows the distribution of the differences between correlations in the A- and V-groups (i.e., rA minus rV), computed by the bootstrap methods with 100000 repetitions. The thick black line is the difference between our correlations (rA-rV=1.45), and the thin lines indicate the CI [0.17 1.87].
